# Functionally dark genes and the transcriptomic landscape of sporulation in a model mushroom-forming fungus

**DOI:** 10.64898/2026.06.13.732014

**Authors:** Csenge Földi, Zsolt Merényi, Árpád Csernetics, Botond Hegedüs, Edit Ábrahám, Zhihao Hou, Xiao-Bin Liu, Bálint Balázs, Dorottya Anna Szafián, Zoltán Lipinszki, László Galgóczy, László G. Nagy

## Abstract

Spores are the primary means of fungal reproduction, contributing to genetic diversity, colonization, and adaptation. Although spore formation is a pivotal part of the fungal life cycle, its genetic underpinnings remain poorly known. In this study, we characterize transcriptomic changes from late meiosis to early basidiospore formation in the mushroom-forming fungus *Coprinopsis cinerea*, decipher several cellular processes, and identify novel genes involved in this process. We identify distinct trajectories of gene expression, each of which display different functional signals, corresponding to meiotic and morphogenetic processes and transitions between these. Our analyses identify diverse arrays of fungal cell wall modifying carbohydrate-active enzymes, ferritins, a putative catechol-melanin synthesis pathway, as well as components of the mitotic/meiotic apparatus. We present twelve highly conserved genes with roles specific to sexual sporulation in both budding yeast and *C. cinerea*, indicating deep conservation of the gene networks driving sexual spore formation. Reverse genetics identified three conserved but functionally poorly characterized genes conferring sporeless and spore-poor phenotypes that result from postmeiotic developmental arrests stemming from spore inflation and nuclear migration problems. Overall, this study provides novel insight into basidiomycete spore formation and highlights the cornucopia of novel functions encoded by functionally dark genes.

## Introduction

The production of basidiospores is the primary means of sexual reproduction in basidiomycete fungi, contributing to genetic diversity, colonization, and adaptation to new environments. Spores of most Basidiomycota are produced on or within fruiting bodies (sporocarps, basidiomes), and are released forcibly *via* ballistospory^1^. A single fruiting body can discharge over a billion spores per day, globally amounting to an estimated 50 million tons of spores annually. This mass of spores has diverse impact on the ecosystem and can even influence weather patterns by attracting water droplets and promoting cloud formation^2^.

While spore production is the most important stage in the life cycle of fungi, the large quantity of spores in industrial mushroom cultivation presents various challenges. Spores can clog ventilation systems, act as vectors for diseases, potentially reducing the yield and quality of harvest. Spores can be harmful to the health of workers^3,4^. After harvesting, mushrooms continue to release spores, which negatively impacts their marketability^5^. Exotic or hybrid mushrooms may unintentionally escape cultivation, and can become invasive or disrupt ecosystems^6^, as demonstrated for the Asian golden oyster mushroom (*Pleurotus citrinopileatus*) in the USA^7^. Sporeless industrial strains offer a potential solution to these problems, yet to date few non-sporulating strains are available for commercial cultivation^8–10^. The sporeless *Pleurotus ostreatus* strain SPOPPO was developed via introgression breeding^11^, taking advantage of a disrupted *msh4* gene which results in the arrest of the first meiotic division and sporelessness^8^.

To facilitate the development of sporeless strains, a more detailed understanding of molecular mechanisms underlying sporogenesis is essential. Spore formation is usually divided into meiotic and postmeiotic stages. The molecular mechanism of meiosis has been well characterized across Dikarya, from yeasts^12^ to filamentous asco-^13,14^ and basidiomycetes^15–18^. Among basidiomycetes, the process of meiosis was most extensively studied in *Coprinopsis cinerea*^15,16,18–20^, which is particularly well-suited for studies of sporogenesis due to its genetic tractability, well-characterized life cycle^21^, and because 60-85% of the approximately 10^7^-10^8^ basidia in a single fruiting body develop in synchrony^21–23^. As a result of classical mutagenesis and reverse genetic studies, numerous ‘white-cap’ *C. cinerea* mutant strains have been described^19,24–26^. These include mutants with meiotic and post-meiotic gene defects^25^ and hence do not produce the black spores characteristic of this species. White cap mutants were later complemented by results of genome and transcriptome studies which also revealed meiotic and post-meiotic genes involved in spore formation^16,27,28^. Whereas the core set of meiotic genes is relatively well established, postmeiotic genes remain less well defined. The conserved transcription factor *Srr1* was shown to be a key regulator of postmeiotic events in *C. cinerea*^27^. Kobukata et al. (2024) identified post-meiotic genes in *P. ostreatus*, and disruption of two of these (*pcl1* and *cro6c*) led to sporeless phenotypes^28^. Despite these advances, much of the molecular and cellular machinery underlying sporulation remains unknown.

A unique group of genes that transcriptome studies consistently bring up are genes of unknown function that encode proteins without any (automatically assignable) annotation terms or conserved domains. Many of these genes encode so-called functionally dark proteins, whose sequences are conserved but lack recognizable or predictable functions^29^. Functionally dark genes are common in fungal genomes and may be a rich source of novel functions and structural elements. A recent comparative transcriptomic analysis revealed more than three hundred families of functionally dark genes, which are upregulated during agaricomycete fruiting body development^30^. One of these (*snb1*) was later shown to be required for proper cell differentiation^31^. Given that the molecular basis of sporogenesis remains only partially resolved, functionally dark genes are especially attractive candidates for discovering new elements of this process.

Previous studies using transcriptome analysis have primarily focused on meiosis^16^, or on a broad timescale^28,32–34^, which did not provide a high-resolution view of postmeiotic processes. In this study we chart the transcriptomic landscape from the end of meiosis to the appearance of sterigmata until spore formation. We had two main objectives: first, to explore post-meiotic processes using a time-series transcriptomic approach; and second, identify and characterize functionally dark genes with the aim of uncovering novel ones related to sporulation.

## Results

### RNA-Seq of postmeiotic sporulation identifies 8 expression trajectories

To investigate post-meiotic steps of sporulation, RNA sequencing (RNA-Seq) was performed on gill samples collected at three consecutive developmental stages (Figure 1/a). Sampling was carried out at the 7^th^, 8^th^, and 9^th^ hour after the start the light phase, corresponding to the onset of meiosis II, the emergence of sterigmata, and the formation of spore initials, hereafter referred to as Stage I, Stage II, and Stage III, respectively (Figure 1/a).Multidimensional scaling (MDS) analysis indicated a close clustering of replicates (Figure 1/b). Differential expression analysis (fold-change(FC)>2, false discovery rate (FDR) ≤0.05) identified a total of 4,475 differentially expressed genes (DEGs) (<u>Table S</u>1). The number of DEGs in each pairwise comparison is shown on Figure 1/a. Of the 4,475 DEGs 1,303 encode proteins without any conserved (Pfam/Interpro) domains, i.e. they are unannotated genes/proteins and 122 that encode proteins that contain domains of unknown function (DUFs).

**Figure 1.**
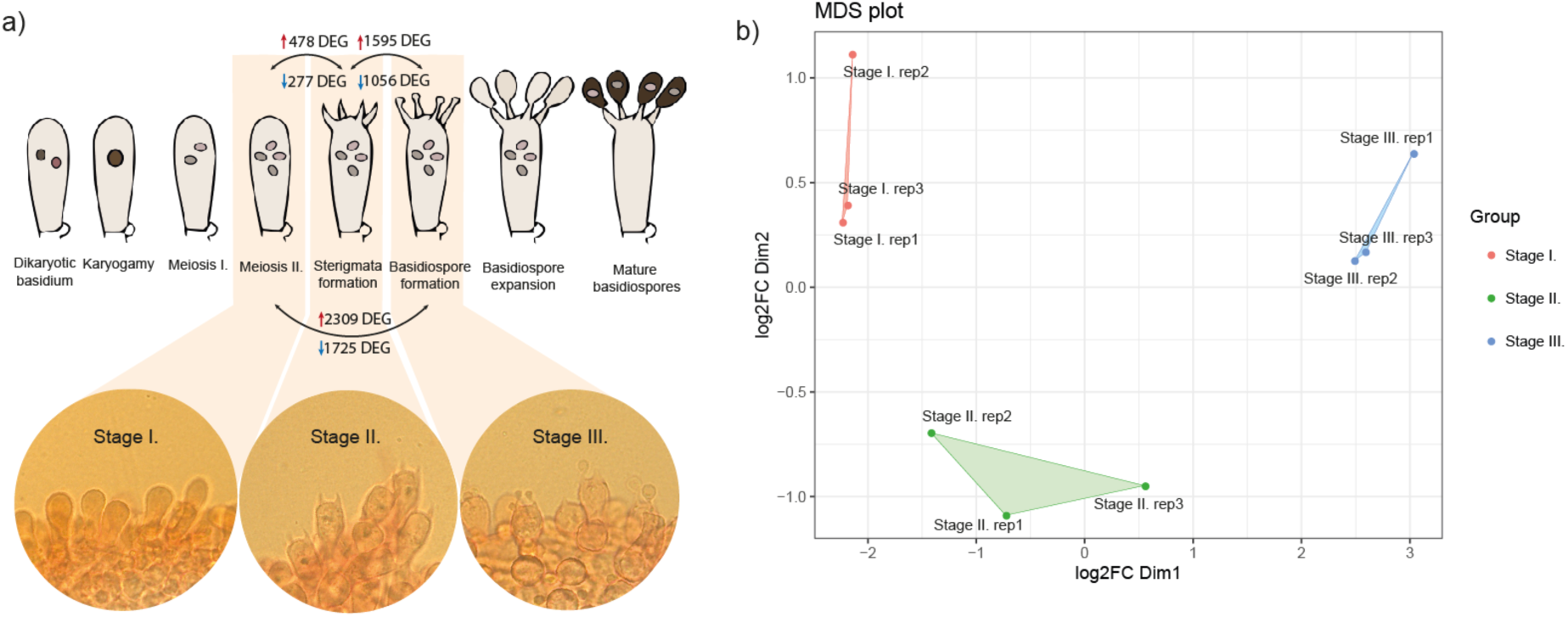
Outline of the sampling strategy for RNA-Seq (a) and the grouping of biological replicates from the three sampled developmental stages (b). Bottom row of panel a shows microscope images of the three stages chosen for RNA-Seq.

DEGs were grouped into clusters using soft (fuzzy) *c*-means clustering^35^. Unlike hard clustering methods, fuzzy clustering assigns a membership value to each DEG, reflecting the degree of association between a gene and a particular expression profile. The optimal number of clusters was determined using the inertia drop and elbow methods^35^, both indicating five clusters. Of these main clusters, three exhibited increasing and two decreasing trajectories, hereafter referred to as clusters I1–I3 and D1–D2, respectively (Figure 2/a). A subset of DEGs with weak cluster membership value (*m* < 0.35, 317 DEGs, 7.08%) was further divided into three subclusters (S1–S3), using the same method. Clusters S1 and S2 displayed expression peaks at Stage II, whereas S3 showed a decline at Stage II (Figure 2/a). Collectively, thus, our analyses arranged DEGs into eight clusters, each having different expression trajectories, capturing the diversity of expression patterns of DEGs across the three developmental stages of sporulation.

**Figure 2.**
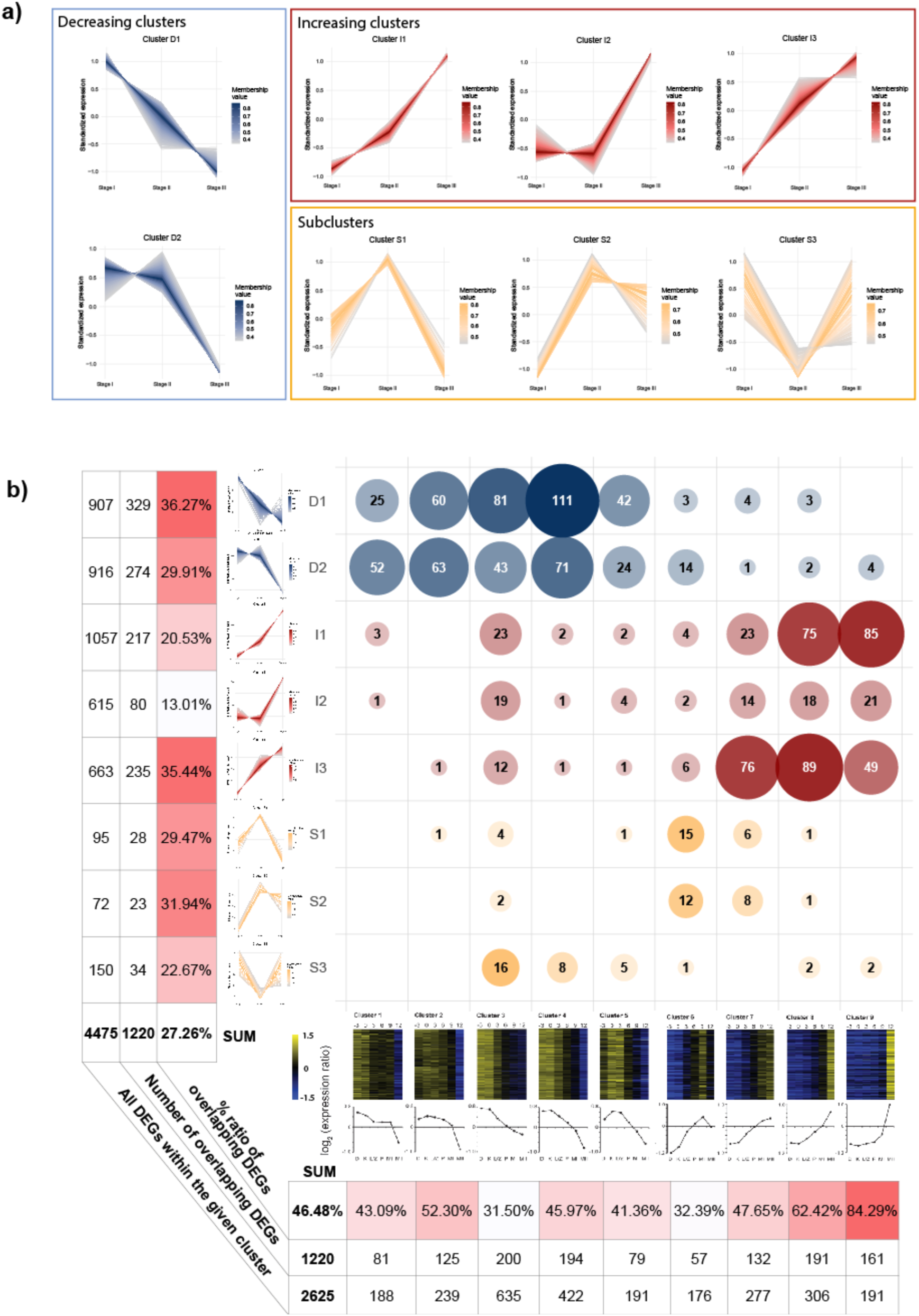
Fuzzy c-means clustering and correspondence with microarray data. a) two decreasing, three increasing and three subclusters that describe the expression trajectories of the 4,475 DEGs in our data. Each individual line in the plots corresponds to an expression trajectory of a gene, colored by membership value. b) correspondence between published microarray data^16^ (columns) and our RNA-Seq data (rows) for meiosis and post-meiotic steps in C. cinerea. Numbers in bubbles correspond to the number of DEGs that overlap between the given pairwise combination of clusters. See also <u>Table S”Burns”</u>.

We compared our results to a previous time-course microarray study, which examined the meiotic gene expression program of *C. cinerea* Okoyama 7 from karyogamy to the end of meiosis II ^16^. Despite methodological differences and the absence of Okayama IDs for a subset of genes, nearly half (46.48%) of the DEGs overlapped between the two studies. Comparison of the two datasets reveals a clear correspondence between temporal expression patterns. DEGs upregulated during early meiosis in Burns *et* al. (meiotic clusters 1–5) overlap mainly with our decreasing clusters (D1, D2), while those upregulated in late meiosis (clusters 7–9) correspond to the increasing clusters (I1–I3). Subclusters S1 and S2, which peak at sterigmata formation, overlap with clusters 6–7 identified by Burns et al, whereas S3, characterized by a decline at sterigmata formation, overlaps with clusters 3–5 (Figure 2/b) (<u>Table S</u>2). This suggests that S3 genes are first upregulated during early meiosis, transiently repressed, and subsequently reactivated post-meiotically—an expression trajectory not captured in the dataset of Burns et al.. Taken together, these results indicate good correspondence between the microarray and our RNA-Seq data, with the latter providing a higher resolution.

### Functional characterization of gene expression clusters

To elucidate the biological processes contained within the eight gene clusters, we integrated Gene Ontology (GO) enrichment analysis (<u>Figure S1GO</u>, <u>Table S</u>3), a list of 721 genes with literature-reported roles, or clear orthology to a yeast counterpart^30^ (hereafter ‘genes with known function’, Table S DEGS/b) and a curated list of 566 carbohydrate-active enzymes (CAZymes)^36^ (<u>Table S</u>4). Overall, a total of 1,256 DEGs were associated with GO terms showing significant enrichment. A comprehensive list of enriched GO terms is provided in <u>Table S</u>3. Cross-comparison with genes with known function identified 287 genes in the 8 clusters (Table S1/b). Hereafter, we functionally characterize each cluster based on GO and genes with known functions; however, providing a comprehensive overview is beyond the scope of this study and therefore additional details are provided in the supplementary materials.

### Decreasing clusters (D1 and D2)

Both decreasing clusters show a general downregulation of gene expression over the course of sporogenesis, albeit with distinct timing and functional associations. Specifically, in cluster D1 (907 DEGs) expression peaks at Stage I, coinciding with meiotic events when basidia contain two nuclei and meiosis I is completed^21^. D1 expression is high prior to sterigmata formation and declines thereafter. Genes in cluster D2 (916 DEGs) maintain a high expression until the end of meiosis II and decrease afterwards.

### Cluster D1 (907 DEGs)

Consistent with cytological events, the results of GO-enrichment analysis of cluster D1 highlighted processes central to meiotic progression. Enrichment was observed for DNA replication and meiotic recombination checkpoint signaling, reflecting the requirement for faithful chromosome segregation. Among the meiosis-specific genes, we detected a *recA* gene family member (1000139) involved in recombination and repair^37^; *rad50* (444936), which has a structural role in axial element and synaptonemal complex formation, homolog pairing and meiotic recombination^38^; *hop1* (464859) required for homologous chromosome synapsis and chiasma formation^39^; as well as *Mnd1* (446459) and *Hop2* (238535) which form a stable heterodimer complex involved in chromosome pairing and repair of meiotic double-strand breaks^40,41^. Many DEGs in this cluster are the part of spliceosomal complexes (U6 snRNP, tri-snRNP and U12-type), which facilitate the high-level transcription and processing of RNAs required for meiosis-specific gene expression^42^.

Terms associated with translation and protein synthesis, including ribosomal subunits and initiation factors, highlight the need for protein production. The protein folding machinery such as the prefoldin complex ensures their proper functionality^43^. The upregulation of the signal peptidase complex may play an important role in the secretion of extracellular enzymes and signaling peptides, including the STE14 homolog isoprenylcysteine carboxyl methyltransferase responsible to CAAX cleavages (39881) and its targets, two mating pheromones (1002221 and 1002223). A further notable term “membrane insertase activity” is highly enriched driven by both insertases of *C. cinerea* (489308 and 385864). Membrane insertases are highly conserved proteins responsible for embedding hydrophobic transmembrane domains of proteins into the lipid bilayer^44^. Cluster D1 also contains several ribosomal protein encoding genes, the galectin genes *cgl1-cgl2* (473274, 488611), the pheromone receptor *rcb3* (394943), the fasciclin *fas1* (500405) as well as the putatively defense-related cocaprin *ccp1* (485770) and cospin *pic1* (367957).

### Cluster D2 (916 DEGs)

For genes in cluster D2 GO terms are strongly enriched for genome maintenance and division, DNA replication and repair, cell-cycle regulation, telomere maintenance, and chromatin organization. Factors central to chromosome stability and segregation are overrepresented. The Smc5/6 complex is essential for maintaining genome stability (e.g. *smc3* - 406915), and has multiple roles in DNA repair and homologous recombination-mediated DNA repair of sister chromatids^45–47^. The NDC80 kinetochore protein complex (453581 and 450001) ensures kinetochore-microtubule attachment and also functions in the activation of the spindle assembly checkpoint^48^ essential for proper chromosome separation and metaphase to anaphase transition. More general GO terms linked to cell division, such as “DNA replication”, “5’-3’ RNA polymerase activity”, “chromatin organization” and “homologous recombination-mediated double-strand break” were also enriched. In parallel, enrichment of RNA processing and ribosomal components align with the observation that the expression of genes encoding ribosomal proteins typically increases before meiosis and gradually decreases during later stages of sporulation^16,49^.

Several signaling components of the mating pheromone pathway were detected, which map into clusters D1/D2. The expression of mating pheromones peak in cluster D1, whereas mating receptors are represented in D2 by two STE3-like GPCRs: one non-mating-type receptor (465723) and one mating-type receptor (356637). Pheromone receptors play a role in the recognition of compatible monokaryons, maintaining dikaryosis, and regulate the post-fusion events in the formation and maintenance of the dikaryon^50,51^. The upregulation of a STE3 mating-type pheromone receptor within young peridioles has been also described in *Pisolithus microcarpus*^32^. The role of non-mating-type pheromone receptors in higher fungi remains unclear^52^, but their presence may be necessary for meiosis and sporulation, as it was observed in *Cryptococcus neoformans* or *Fusarium graminearum*, where non-mating-type receptors have been proven essential for meiosis and sporulation^53,54^. Therefore, it is conceivable that non-mating-type receptors in *C. cinerea* may play a similar role. Cluster D2 also contains the chitinase *ChiEN4* (358869) and the cyclopropane fatty acid synthase-1 *cfs1* (447925) as well as the sumoylation factor *Cc.sumo1*^55^ (446185).

### Increasing clusters (I1-I3)

Genes showing increasing expression trends were grouped into three clusters (I1-I3) (Figure 2/a). Cluster I1 (1,057 DEGs) and I3 (663 DEGs) show monotonously increasing expression, whereas I2 (615 DEGs) includes genes whose expression is low in the first two stages, and increase rapidly in stage III. It is noteworthy that most differentially expressed CAZymes are found in increasing clusters, but not in decreasing or subclusters. Of the 556 CAZymes encoded in the *C. cinerea* genome, 227 are differentially expressed. Of these 168 are in increasing clusters, 44 in decreasing and 14 in subclusters. The concentration of CAZymes in increasing clusters might be a consequence of the activation of morphogenetic events in stages II and III, which involve sterigmata and spore formation and cell wall remodeling. This is also reflected in the enrichment of the “carbohydrate metabolic process” GO term in increasing clusters. Given the importance of CAZymes to fungal morphogenesis, we pay special attention to this enzyme class in the following descriptions of increasing clusters.

### Cluster I1 (1057 DEGs)

I1 is the largest cluster, showing a gradual increase in expression across the three sampled stages. GO enrichment revealed terms related to carbohydrate metabolism, transmembrane transport, cell surface receptor signaling, extracellular region, vacuolar and cell membrane, amino acid metabolism, among others (<u>Table S</u>3).

Transport- and membrane-associated terms, especially those linked to the vacuolar membrane could be related to the formation of a vacuole at the base of the basidium which progressively expands as the cytoplasm and other organelles migrate into the developing spores. By the time the spores mature, the vacuole occupies nearly the entire basidium^56^. We found that the enrichment of the GO term cell surface receptor signaling corresponded to the upregulation of three G-protein α-subunits (IDs: 354280, 363001, 377794) and one β-subunit (IDs: 503715) and a Git3 glucose receptor (426382).

Carbohydrate metabolism, particularly (1→6)-beta-D-glucan biosynthetic process, cellulose- and chitin binding showed a strong signal in this cluster. Corresponding DEGs may participate in cell wall synthesis (e.g., chitin synthase, β-1,6-glucan synthase) and modification (e.g., chitinases, chitin deacetylases)^30,57^. We identified 30 glycoside hydrolases in cluster I1, which include glucanases (e.g. GH16) and chitinases (e.g. GH18) but also other activities, and, along with two putative expansins, may be involved in cell wall loosening and modification. Chitinases include *ChiEn2* (90984), *ChiEn3* (470416), *ChiE1* (543586) and *ChiE2* (520359). The latter was shown to be related to sporulation^27^.

This cluster also includes members of the glycoside hydrolase 13 family, showing α-amylase-like activity, with all *C. cinerea* members differentially expressed. These have been previously implicated in structural rearrangement of the fungal cell wall through α-glucan synthesis and modification^57,58^. Two chitin deacethylases (CE4) were found in this cluster, these convert chitin to the more recalcitrant chitosan that may form crosslinked matrices with other cell wall polymers^59^. Further seven DEGs encoding CBM1 domain proteins which, beyond cellulose, may be able to bind fungal cell wall polymers^30^. The five ricin-B lectins (CBM13) may have functions in cell attachment or defense. We further found ten glycosyl transferases, of which 4 belong to GT2 chitin synthases and may be involved in producing new cell wall materials.

Among experimentally characterized genes, the carbon catabolite repressor *cre1* (466792), the putative light sensor *dst2* (361574) and the photolyase/cryptochrome *phr1* (492962) were also assigned to this cluster.

### Cluster I2 cluster (615 DEGs)

Cluster I2 includes DEGs upregulated during spore initiation (Stage III) (Figure 2/a). DEGs in this cluster are associated with 14 enriched GO terms, among which the most significant include N-acetylglucosamine catabolic process, carbohydrate metabolism, and extracellular space. Notably, this cluster also contains many CAZymes, including 16 AAs, seven carbohydrate binding modules, three CE4 chitin deacethylases, one expansin, 15 GHs and three glycoside transferases. The low number of GT genes (compared to cluster I1) suggests that emphasis shifts from synthesis to modification of the cell wall. Auxiliary activities (AA) include three AA1 multicopper oxidases (e.g. laccases) and six lytic polysaccharide monooxygenases (AA9). The AA1 family included the laccases *lcc3* and *lcc8*, which are noteworthy from the perspective of melanin production, with which laccases have been associated. It should be noted that AA enzyme-encoding genes were missing in cluster I1 but are abundant in I2.

This cluster further included two septins (415573, 455217), which show high activity when spores emerge at the sterigmata tips. As conserved GTP-binding proteins, they might contribute to spore formation by directing membrane morphogenesis and wall assembly. The cluster also includes the recently described RNA binding protein gene *ort2*^60^ and the fungalysin *mep2*^61^.

### Cluster I3 (663 DEGs)

Genes in Cluster I3 show monotonous increase in expression from Stage I, and reach highest expression in Stage III. GO enrichment in cluster I3 highlights 23 GO terms, including gluconeogenesis, purine and amino acid biosynthesis, redox cofactor metabolism and cell polarity establishment. Gluconeogenesis is one of the most significantly enriched terms. In budding yeast, glycogen breakdown and gluconeogenesis proceed in parallel during mid-meiosis, channeling carbon from both glycogen and acetate into spore wall synthesis^62^. Although the extent to which yeast findings can be extrapolated to *C. cinerea* is unknown, there are noteworthy parallels with DEGs identified in our data. These include fructose-2,6-bisphosphatase, which depletes the allosteric activator of phosphofructokinase, thereby enforcing a unidirectional gluconeogenic flux that supports cell or spore wall assembly. Additional contributors are fructose-aldolases (546961, 408165) and fructose-1,6-bisphosphatase (436876), which further sustain gluconeogenic flux. In addition to these, we identified an acetate transporter (441597) in cluster I3, which may also contribute to spore wall assembly.

Of the experimentally characterized *C. cinerea* genes, cluster I3 contains *Eln2, Eln3,* the ortholog of the *Schizophyllum commune* transcription factor *hom2*, the fungalysin *mep3*, the light responsive gene *nod1*, the putative chromatin-associated *ich1* and the ortholog of *pcl1* of *P. ostreatus*, which confers a sporeless phenotype when disrupted^28^. It also includes *psr2* and *psr3*, two unannotated genes characterized below. Of the CAZyme encoding genes, cluster I3 contains eight AA genes, four CBMs, one expansin, twelve GH and seven GT (2 chitin synthases) genes. Interestingly, AA-s included glucose-methanol-choline oxidoreductases and glyoxal oxidases, but no laccases or AA9 like in cluster I2.

### Subclusters (S1-S3)

Subclusters were obtained by reclustering genes with low membership value from main clusters. As a result, these clusters are smaller and contain 72-150 DEGs. Clusters S1 and S2 both include DEGs that show an expression peak at Stage II, corresponding to the end of meiosis and the emergence of sterigmata. The main distinction between clusters S1 and S2 lies in post-Stage II dynamics: S1 genes peak sharply at Stage II before returning to or below baseline at Stage I levels, whereas S2 genes also peak at Stage II but remain elevated thereafter. Cluster S3 genes show a transient decrease at Stage II, indicating temporary suppression during this phase. Since each of the subclusters contains a relatively small number of DEGs, the number of enriched GO terms are also lower than in other clusters.

#### Cluster S1 (95 DEGs)

DEGs in this cluster are associated with six enriched GO terms, including phagocytic vesicle, mitotic chromosome condensation and anatomical structure morphogenesis. We found no experimentally characterized gene in this cluster.

#### Cluster S2 (72 DEGs)

We identified an enrichment of G protein-coupled receptor (GPCR) signaling related terms, which correspond to two G-protein alpha subunits (367330 and 465055) belonging to this cluster. The cluster also includes three transcription factors (451915, 366541 and 543308). One of these is SRR1, recently described as a post-meiotic regulator of spore formation in *C. cinerea*^63^. This cluster also includes *psr1*, which is functionally characterized below.

#### Cluster S3 (150 DEGs)

This cluster shows strong enrichment for five GO terms, including heme binding and lipid metabolic processes. Among the enriched genes are those encoding cytochrome P450 enzymes, heme peroxidases, fatty acid desaturases, and fungal lipases. In addition, genes involved in reciprocal meiotic recombination are also represented, including the SPO11 orthologue topoisomerase. Interestingly, this cluster includes the light receptor *C. cinerea wc2,* the lectin *ccl1*, the morphogenetic gene *snb1* and the putative epigenetic regulator *Cc.tet*^64^.

### Ferritins are highly upregulated during sporulation

Within increasing clusters, we identified a DEG group that is not associated with any GO-term but characterized by very high expression levels (Table S1). The expression of genes encoding ferritins based on conserved domain annotations (51853, 202468, 547137) was strongly upregulated when spores emerge (fold change 288 - 3,684), while their expression remains negligible until sterigmata formation. Two of them (51853, 547137) show undetectable expression levels in the Δ*srr1* sporeless mutant strain^63^, suggesting that spore initiation is a prerequisite for their upregulation. Ferritins are important intracellular iron-storage proteins, sequestering and buffering iron. Ferritin accumulation has also been observed in *Phycomyces* spores^65^, in which iron is released upon germination^66^. David and Easterbrook (1971) proposed that ferritins serve as a mobilizable iron reserve at germination, supporting early iron-dependent processes ^65^. In *Fusarium*, early germination is known to impose iron demands on biosynthesis and energy metabolism^67^. Based on our data a similar role is plausible in *C. cinerea*.

### *C. cinerea* may utilize the catechol-melanin pathway in spores

Our RNA-Seq data offer an opportunity to investigate the genetic basis of melanin production. Agaricomycete genomes encode multiple melanin biosynthetic pathways^68^, all of which start from chorismate, which also serves as a central precursor for the biosynthesis of various aromatic compounds^69^, including aromatic amino acids^68^. In cluster I1 a shikimate dehydrogenase gene (543926) showed a 23-fold, while a chorismate synthase gene (265487) showed a 55-fold increase in expression at Stage III compared to Stage I. This overlaps with observations made recently on *C. cinerea* gill transcriptome data^63^ and may be indicative of active melanin synthesis pathways. Additionally, the shikimate dehydrogenase and chorismate synthase gene (543926 and 265487) were downregulated in a sporulation-deficient transcription factor mutant^63^.

We therefore checked components of the four melanin synthesis pathways described in *Agaricus bisporus*, based on orthology with *C. cinerea* genes. Within the examined time window, genes associated with the L-DOPA melanin pathway, described as the predominant melanin biosynthesis route in Agaricomycetes^68,70,71^, did not show marked upregulation (Table S5). In contrast, genes encoding enzymes linked to the catechol-melanin pathway were induced, such as a phenylalanine ammonia-lyase (361541, FC_(III/I)_: 4.88), a 4-coumarate-CoA ligase (191140, FC_(III/I)_: 18.89), and a catechol dioxygenase (427648, FC_(III/I)_: 3.0), suggesting their involvement in spore pigmentation. Previous RNA-Seq data partially agree with this contention: throughout the development of *C. cinerea*, the 4-coumarate-CoA ligase and catechol dioxygenase genes have highest expression in young fruiting body gills, where sporulation starts^72^.

The catechol-melanin pathway has previously been described in few Basidiomycetes, including *Rhizoctonia solani*^73^ and *Ustilago maydis*^74^. Although chorismate is presumed to act as a precursor for catechol-type melanin biosynthesis^75^, the exact enzymatic steps of this pathway remain incompletely characterized in fungi. Nevertheless, our data suggests that, despite the genomic presence of multiple melanin biosynthetic genes in *C. cinerea*, catechol-derived melanin may be the dominant form during spore development. Nevertheless, it should be noted that alternative melanin synthesis pathways may also exist, as recently demonstrated in *U. maydis*^76^.

### Ancient genes dominate the sporogenesis transcriptome

To uncover the evolutionary/temporal dynamics of genes active during sporulation we assessed the age distribution of DEGs in each of the clusters using phylostratigraphy (see Methods) (Figure 3). Overall, most genes were either very ancient or young (i.e.*Coprinopsis*-specific). Decreasing clusters (D1/D2) are significantly enriched in ancient genes, consistent with their enrichment in mitotic and meiotic processes. In contrast, increasing clusters showed different or no enrichment patterns. For example, the origins of genes in cluster I1 concentrated in early fungal nodes, indicating potential innovations in postmeiotic sporulation. Genes in cluster I3 were enriched coincident with the origin of Basidiomycota.

**Figure 3.**
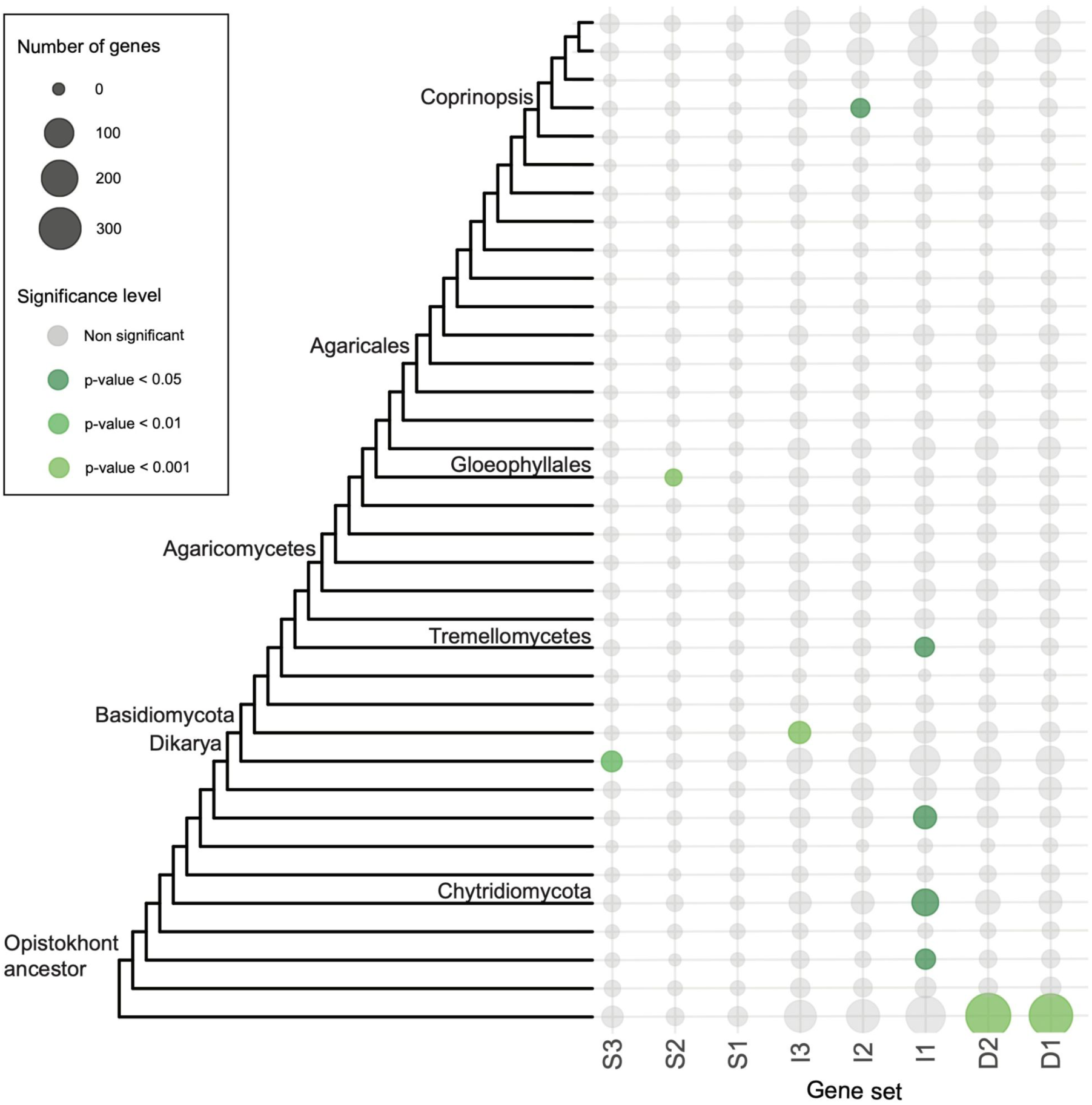
Phylostratigraphic analysis of gene ages in the eight clusters. We mapped the origin of each gene to specific nodes of a phylogeny, based on the most distant species in which clear sequence-based orthologs can be detected. Bubble size and color are proportional to the number of genes mapping to a specific node and significance level, respectively

Given the presence of evolutionarily conserved genes in most clusters, we examined whether our DEGs have clear orthologs in *Saccharomyces cerevisiae*. Although *S. cerevisiae* and *C. cinerea* differ in their sporulation mode, in that the former produces ascospores internally whereas the latter produces basidiospores externally, any clear ortholog with a sporulation-specific role in the two species could indicate a highly conserved genetic program for sexual sporulation. Such patterns clearly exist for meiotic genes (see)^16^, therefore we focused on postmeiotic clusters, in search of deep conservation of spore morphogenesis genes. We identified 71 *S. cerevisiae* genes involved in sexual spore formation (see Methods). Of these 33 had putative orthologs in *C. cinerea* and 19 were differentially expressed in our dataset. These were mostly found in clusters I1-I3 (Table S1/c). Twelve of the 19 proteins have a sporulation-specific function in yeast, whereas the other seven have broader biological roles (e.g. G-proteins, protein kinase C). The twelve genes included inducers of the *S. cerevisiae* meiosis-specific transcription factor IME1 (EMI1, EMI2) sporulation-specific cell wall remodeling chitinases and glucanases and their activator (CTS2, SPR1, SHC1), a sporulation-specific septin (SPR28), an anaphase promoting complex activator required for spore wall assembly (AMA1), a spore-specific water channel (AQY1), a STE20-family GCKIII protein kinase; required for prospore membrane closure (SPS1) as well as poorly known proteins (SPS18, SPS19, SPO75) (Table S1/c). The presence of these genes suggest that orthologs specific for sexual sporulation are conserved in *C. cinerea*, and likely play roles similar to those in *S. cerevisiae*.

We made two surprising observations in this gene set. First, eight genes are in increasing clusters, indicating a post-meiotic role. In contrast to meiotic genes, postmeiotic genes (e.g. involved in spore morphogenesis), are expected to evolve faster, and potentially not overlap between endogenously produced ascospores and exogenously produced basidiospores. Second, this suite of genes includes several regulators and protein kinases (SPS1, SHC1, EMI1-2, AMA1). Such a high conservation of regulatory genes is unexpected and we hypothesize that this reflects their involvement in a highly conserved sexual spore development program that is shared between Asco- and Basidiomycota, dating back to at least the divergence of these phyla, approximately 600 million years ago^77^.

### Conserved, functionally dark genes

It is noteworthy that over half of the DEGs (54.66%) have no associated GO terms, leaving their biological roles uncharacterized and thus absent from the functional categories discussed above. A high proportion of functionally poorly characterized genes has already been noted across studies involving mushroom fruiting bodies, including, for example, morphogenesis and sporulation^31,63^. Poorly characterized but highly conserved genes are especially intriguing, as they may encode completely new functions underlying fundamental molecular processes. We therefore scrutinized such genes based on a previous comparative transcriptome study (1). Of the unannotated and functionally poorly characterized genes of *C. cinerea* of that study, 157 were differentially expressed in our dataset (Figure 4), <u>(Table</u> <u>S6)</u>. These are distributed in our clusters as follows: 62 are in decreasing clusters, 79 in increasing clusters and 16 in subclusters. In the context of data covering the complete developmental process of *C. cinerea*^72^, the expression of the 157 genes typically peak late during development, often in gills (Figure 4), supporting their association with spore formation.

**Figure 4.**
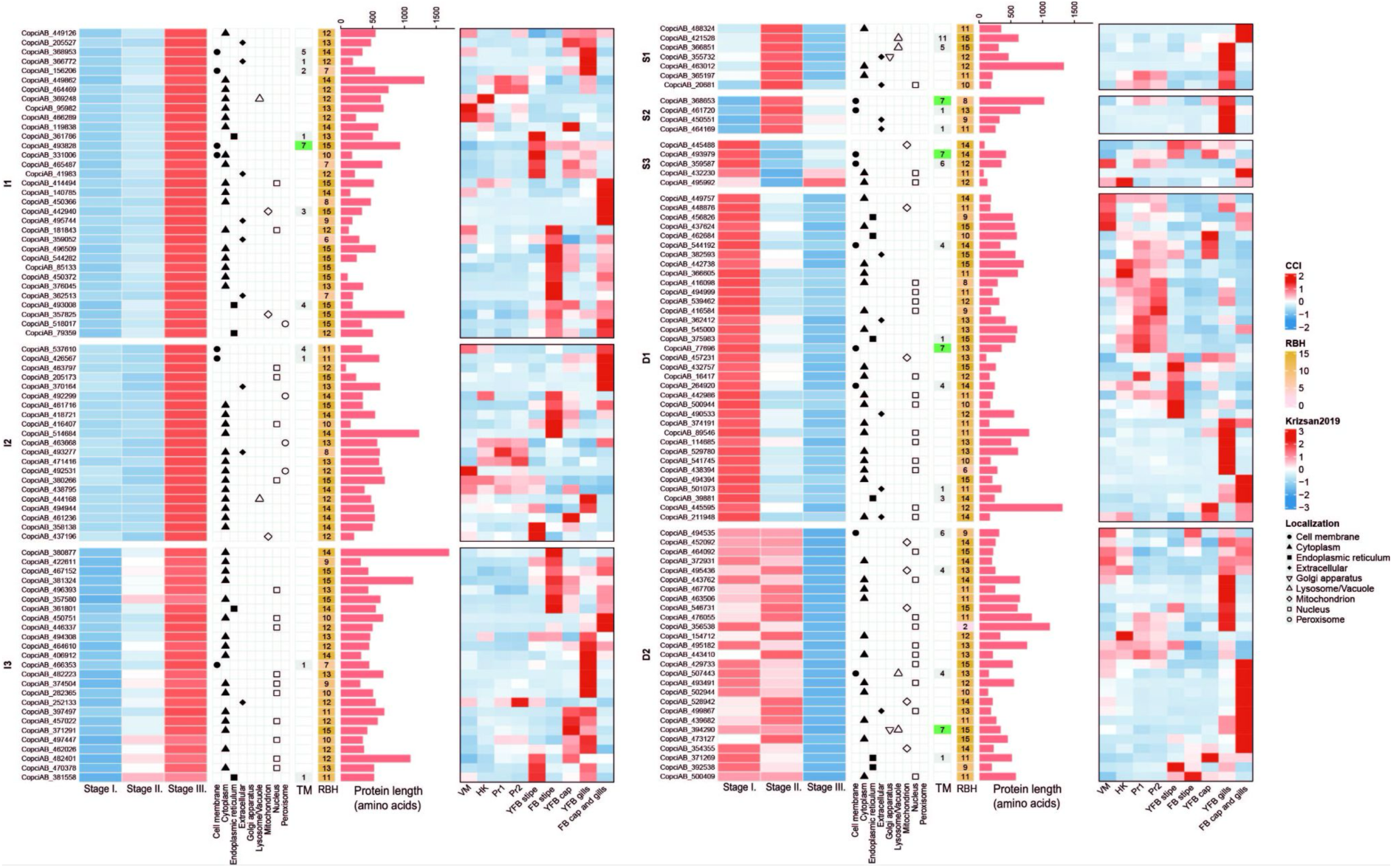
Expression, conservation and properties of conserved, but functionally poorly characterized genes. In the right heatmaps abbreviations are as follows: VM - vegetative mycelium, HK - hyphal knots, Pr1 - primordium 1, Pr2 - primordium 2, YFB - young fruiting body and FB - mature fruiting body. Heatmap legends CCl and Krizsan2019 refer to the left, 3-stage and the right, 8-stage heatmaps, respectively. RBH indicates the conservation of the given gene, in the context of presence/absence of reciprocal best blast hits in 15 species from Nagy et al 2023.

The 157 genes display diverse characteristics in terms of amino acid sequence, predicted localization and transmembrane domains. They are predominantly cytoplasmic (81; 52%) or nuclear (42; 27%), and range in length from 66 to 1,794 amino acids (Figure 4). Twenty-two of the nuclear proteins belong to decreasing clusters, suggesting they may represent hitherto uncharacterized components of the mitotic or meiotic machinery specific to Agaricomycetes. Based on SignalP prediction, 16 proteins are targeted to the cell membrane and another 19 are extracellular. Seven secreted proteins are shorter than 300 amino acids, qualifying them as small secreted proteins, which are often involved in cell-cell communication and fungal development^78^, among other functions.

Overall, we detected a high number of under-characterized conserved genes that may play key roles in basidiospore development.

### Reverse genetic analysis of three conserved functionally dark genes

Three highly conserved, but functionally dark genes were selected for reverse genetic analysis (461720, 482223, 450751). These have an expression peak in the gills during development of *C. cinerea* (ref)^72^, have the highest expression in Stage II or III in our dataset (Table S1), and have orthologs in other fruiting body-forming basidiomycetes, including the industrially relevant *P. ostreatus*. The three selected genes were designated as “putative sporulation related” (psr) genes *psr1* (461720) from cluster S2 and *psr2* (482223) and *psr3* (450751) from cluster I3.

Based on structural predictions on the encoded proteins (see Methods), PSR1 is localized in the cell membrane and contains a transmembrane domain, whereas PSR2 and PSR3 contain a nuclear localization signal. InterPro domain predictions indicated that PSR1 and PSR2 lack any known conserved protein domains, while PSR3 is predicted to contain an integrase zinc-binding domain (IPR041588), for which no more precise function could be assigned.

The three genes were disrupted using an *in vitro* assembled RNP-based CRISPR/Cas9 system^79^ to investigate morphological changes during fruiting body development resulting from their loss. In wild-type fruiting bodies, the gills turn black upon maturation due to the accumulation of dark pigmented spores. In contrast, mature fruiting bodies of the *Δpsr1* and *Δpsr3* mutants remained completely white, whereas those of the *Δpsr2* mutant displayed a light brownish coloration (Figure 5/feno?).

**Figure 5.**
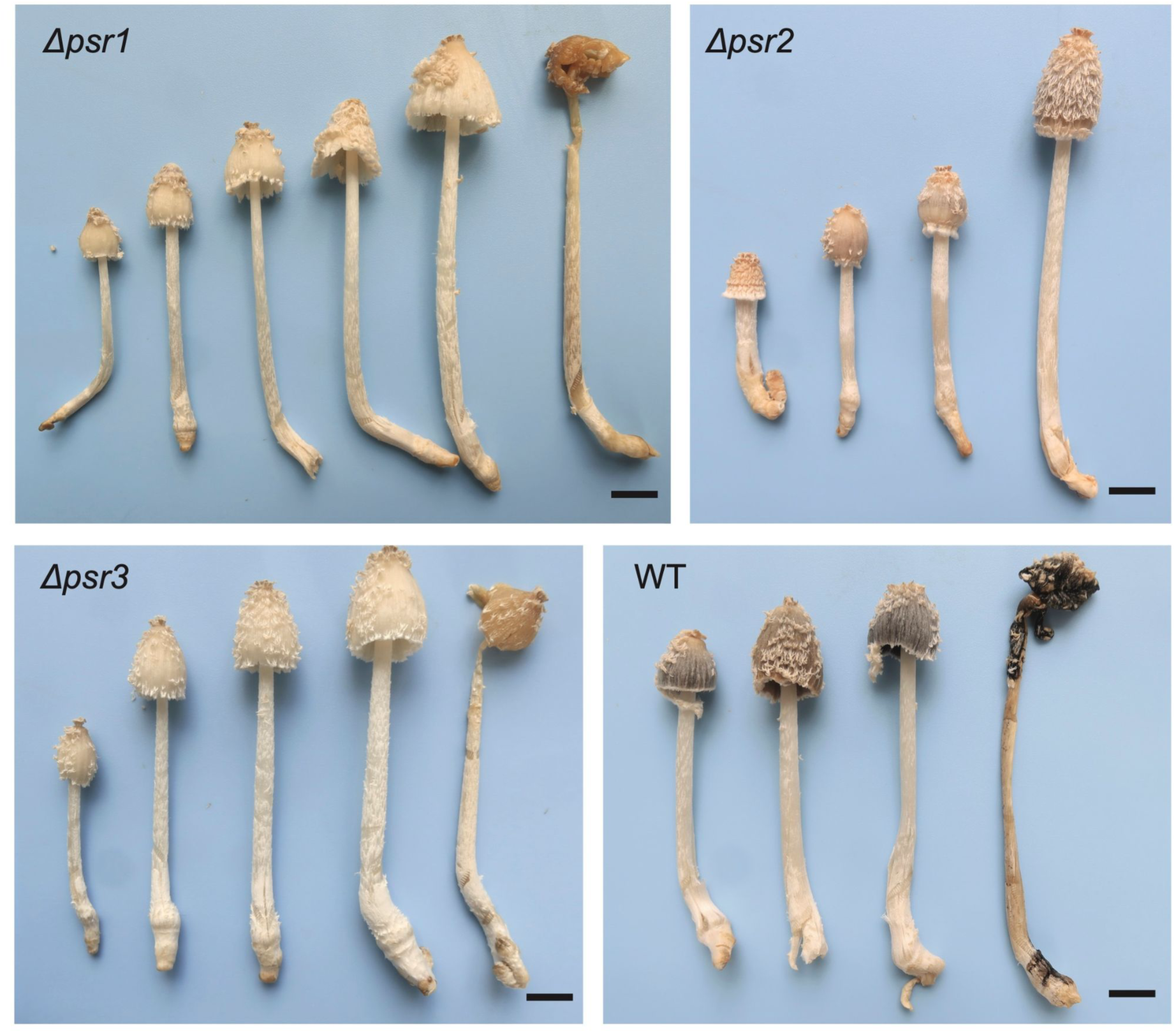
Mutant and wild type fruiting bodies at multiple developmental stages, highlighting the lack of black sporulation in Δpsr1 and Δpsr3, and the spore-poor phenotype of Δpsr2. Scale bar represents 1cm. See supplementary video at https://www.youtube.com/watch?v=ZCY4q4ffbus

We also examined fruiting bodies at the stage when, in the wild type, spore maturation was complete and spores had begun to detach from the sterigmata, but autolysis had not yet started (Figure S3). In wild type, each basidium produced four sterigmata, each bearing an ovoid, dark brown spore (11–12 × 6.5–7 µm, n=), clearly visible without staining. In the *Δpsr1* mutant, most basidia appeared bare, with only a minority bearing sterigmata and swollen spores. In *Δpsr3* swollen spore initials were observed, but pigmentation and detachment were absent. In this mutant, the proportion of basidia with sterigmata and swollen spores was comparable to wild type. The *Δpsr2* mutant produced far fewer spores than wild type; these spores were pale brown, drop-shaped, and smaller in size (7.9–8.6 × 5.5–5.9 µm) (Figure S2 *Δpsr2*). While the wild type strain produced an average of 3.5 million spores per 100 mg of mature cap tissue, the *Δpsr2* strain produced only ∼133,000 spores (3.8 % of wild type).

To assess whether meiosis can be completed by the mutants, we examined basidia using fluorescent microscopy. In all three mutant strains, four nuclei were clearly visible within each basidium, indicating defects in post-meiotic spore formation. In wild-type strains, nuclei migrated into the developing spores. In the *Δpsr1* and *Δpsr3* mutants swollen spores were present on the sterigmata; however, we could not observe the migration of nuclei into the spores. In the *Δpsr2* mutant, we could not observe a clear trend in nuclear migration, but in some basidia all the four nuclei migrated into a single spore (Figure S3).

## Discussion

This study investigated the transcriptomic landscape of spore formation in the model mushroom *C. cinerea,* with an emphasis on postmeiotic processes and genes of poorly understood functions. Our study highlights the complex transcriptional changes during sporogenesis and underscores the challenge posed by a scarcity of functional knowledge in Agaricomycetes. Nevertheless, our data outline distinct gene expression trajectories and functional categories associated with the progression of spore formation.

While most earlier studies focused on meiosis, we capture post-meiotic dynamics, including sterigmata formation and spore inflation. These stages possess distinct transcriptional programs which are enriched in genes related to morphogenetic processes, cell wall remodeling, carbohydrate metabolism, and iron-storage. The striking induction of ferritin-encoding genes at the onset of spore emergence illustrates how RNA-Seq can pinpoint candidate genes with specific developmental roles that were previously overlooked. Given the importance of iron storage and mobilization during fungal germination^66^, these findings open new avenues for exploring how metal homeostasis underpins spore viability. Another unexpected observation was that our data point in the direction of the catechol-melanin pathway being responsible for melanization of basidiospores. This is a rarely discussed melanin-synthesis pathway in Agaricomycetes^80^; our observations indicate that a broader diversity of melanin-synthesis pathways may need to be considered in future studies.

Our dataset is also a valuable source for mining novel gene functions that currently reside in unannotated gene families - a ‘dark matter’ in agaricomycete genomes. We identified over 150 functionally completely uncharacterized, but conserved genes that showed dynamic expression during sporulation. Reverse genetic analyses of three of them showed that such genes can comprise novel, hitherto unrecognized functionalities. The sporulation defects observed in gene disruptants also indicate that the genes identified here are not only differentially expressed but also functionally critical, thus representing promising targets for developing sporeless strains of cultivated fungi. This is particularly relevant for industrial production, where spore release remains a significant ecological and health concern^28^. In this context, although *C. cinerea* as a model provides unique advantages due to its synchronous progression of meiosis^21^, extending these analyses to industrially important species such as *P. ostreatus* will be essential to translate basic discoveries into applications.

In conclusion, our study fills a gap by providing a new omics dataset on sporogenesis and demonstrates how integrating transcriptomics with reverse genetics can unravel both known and novel components of spore formation. Beyond expanding our basic understanding of fungal development, these insights hold practical implications for the development of sporeless industrial strains.

## Methods

### Strains and culture media

The p-aminobenzoic acid (PABA) auxotrophic homokaryotic *C. cinerea* AmutBmut1 pab1-1 #326 strain was cultured and maintained in a YMG medium. The composition of media and buffers used in the transformation (YMG, minimal medium, regeneration medium, top agar MM- and MMC buffers, and PEG/CaCl2 solution) followed the description of Dörnte and Kües (2012) (100). To obtain oidia, *C. cinerea* was cultured on YMG at 37°C under continuous light for 6 days.

### RNA-Seq analysis

For RNA-Seq samples were taken from the central gill region of *C. cinerea* fruiting bodies. Three developmental stages were collected: after seven hours of light exposure (late meiosis I transitioning to meiosis II, prior to sterigmata formation), after eight hours (completion of meiosis II, with basidia containing four nuclei and sterigmata visible), and after nine hours (initiation of spore inflation on sterigmata)^21,23^. Each stage was sampled in triplicates and stored at –80 °C until RNA isolation.

RNA extraction and library preparation were performed as described by Földi *et* al. (2024) with minor modifications. Briefly, samples were washed with diethyl pyrocarbonate-treated water, homogenized in liquid nitrogen, and total RNA was isolated using the Quick-RNA Mini-Prep Kit (Zymo Research). RNA integrity and quantity were assessed by gel electrophoresis and Nanodrop spectrophotometer. Strand-specific cDNA libraries were prepared from poly(A)-enriched RNA using the Illumina TruSeq Stranded kit and sequenced on the Illumina Novaseq 6000 platform (S4 PE150) at Novogene (UK), yielding 39.7–51.9 million paired-end reads per sample.

Read preprocessing was performed with bbduk.sh (BBMap suite) by trimming adaptors, ambiguous bases, and low-quality regions (trimq = 30, minlen = 50). Reads were aligned to the reference genome (https://mycocosm.jgi.doe.gov/Copcin2/Copcin2.home.html)^36^ using two-pass STAR alignment^81^, following the same settings as in the previous study of Merényi et al.^82^. On average, 13.8–18.3 million fragments per sample were assigned to features. Expression levels were quantified as Fragments Per Kilobase of transcript per Million mapped reads (FPKM), and genes were considered differentially expressed if they showed a log fold-change (LogFC) greater than 1 or less than –1 between any two stages, with an adjusted *p*-value ≤ 0.05 (Benjamini-Hochberg correction). Genes with FPKM < 5 in all samples were excluded.

### mFuzz clustering

Differentially expressed genes (DEGs) were clustered according to their temporal expression dynamics using the *mFuzz* R package (version: 2.68.0), which implements soft (fuzzy) *c*-means clustering (47). Prior to clustering, FPKM values were log₂-transformed and subsequently standardized to remove systematic differences in scale between genes. The optimal number of clusters was determined using the inertia drop and elbow methods. Following this, the fuzzifier parameter (*m*) was estimated with the *mestimate* function and applied in the clustering procedure. Genes were assigned to clusters with membership values between 0 and 1, reflecting the strength of association with a given expression profile. DEGs with weak membership values (*m* < 0.35) in the main clusters were subjected to an additional round of clustering, which yielded three subclusters.

### Functional annotations

Conserved protein domains were identified using InterProScan (version 5.61-93.0)^83^. Gene Ontology (GO) enrichment analysis was performed with the R package *topGO* (version 2.44.0)^84^. Enrichments were discarded if only one gene of a particular class were identified. Enrichments with a Fisher’s exact score >0.05 were also discarded. The cellular localization and the presence of signal sequences were predicted using DeepLoc-2.1. (105) (Odum, 2024). The number of transmembrane domains was predicted by DeepTMHMM (version: 1.0.24.)^85^.

### Analysis of orthology between S. cerevisiae and C. cinerea

To identify ascospore-related genes, we collected genes with the GO terms ‘sporulation resulting in formation of a cellular spore’, ‘ascospore formation’, ‘prospore septin filament array’ and ‘regulation of sporulation resulting in formation of a cellular spore’ from yeastgenome.org^86^. We then identified orthologs of these genes in *C. cinerea* based on blast similarity, by classifying yeast proteins into four groups: proteins with clear reciprocal best hits in *C. cinerea*; proteins with one *C. cinerea* hit that is 10 orders of magnitude better than the next hit (referred to as best blast hit); proteins showing no clear orthologs, or no blast hits at all in *C. cinerea* (e<10^-5^). Only the first two categories were further considered.

### Phylostratigraphy

To evaluate whether genes in DEG clusters are enriched in specific evolutionary ages (phylostrata) we employed a phylostratigraphic approach using a dataset of 655 genomes (G655; Virágh *et* al.) representing all Holomycota phyla derived from the Opisthokont ancestor. Phylostratum assignments were made through a reciprocal best-hit (RBH) method. We performed bidirectional homology searches between the CopciAB 2.0 genome^36^ and the 655 genomes using MMseqs2 (R15-6f462) at a sensitivity of 5.7 for three iterations, retrieving up to 5,000 target sequences (E-value < 0.0001). A bidirectional coverage filter of 40% was applied to the results. We retained only RBH connections, where gene A from species 1 matches gene B from species 2, and *vice versa*. For each CopciAB gene, its “birth node” in the 655 species tree was calculated as the most recent common ancestor of all species with an RBH connection. Genes without RBH connections were deemed strain-specific and assigned the most recent age. Over-representation analysis was conducted using the ora function of the mulea package (HIV)^87^, testing input DEG cluster gene sets, against the background distribution of all gene ages. Significance was assessed using empirical false discovery rate (eFDR) correction based on 10,000 permutations.

### CRISPR/Cas9 based gene disruption

Single-guide RNAs (sgRNAs) were designed using *CRISPOR* (Concordet & Haeussler, 2018) based on the exon regions of the *psr1-2-3* genes (protein IDs: 461720, 482223, 450751). The *C. cinerea* reference genome (GeneModels_FrozenGeneCatalog_20160912.fasta) was retrieved from JGI MycoCosm^88^. sgRNA candidates followed the (N)20-NGG protospacer-PAM rule, with selection criteria minimizing mismatches across the genome, and targeted the first exons of the genes of interest. gRNAs are listed in <u>Table S”Primers”</u>. crRNAs and tracrRNAs were synthesized by IDT (Integrated DNA Technologies) and annealed by mixing 1.2 µL of 20 µM crRNA and 1.2 µL of 20 µM tracrRNA in 9.6 µL duplex buffer, incubating at 95 °C for 5 minutes, then cooling to room temperature. To assemble the ribonucleoprotein (RNP) complex, 0.5 µL duplex buffer, 1.5 µL Cas9 buffer (as described by Liang et al., 2018^89^), and 1 µg TrueCut HiFi Cas9 (Invitrogen, A50575) were added. The reaction was incubated at 37 °C for 15 minutes. The prepared RNPs were stored on ice and 15 µL was used per protoplast transformation.

Circular repair plasmids were based on a pUC19 backbone and carried the *pab1* gene for positive selection, flanked by 1 kb-long homologous arms targeting the genes of interest, as described by Dörnte and Kües (2012)^90^. Homology arms were PCR-amplified using overlapping Gibson primers starting 10 bp upstream of the predicted Cas9 cleavage site. The *pab1* cassette, including its native promoter and terminator, was amplified from the pMA412 vector^91^ and inserted between the flanking arms. All fragments were purified using the QIAquick PCR Purification Kit (Qiagen). Assembly of the repair plasmid was performed with an enhanced Gibson Assembly Mix according to Rabe and Cepko (2020)^92^. All PCRs were performed with Phusion Plus High-Fidelity DNA Polymerase (Thermo Scientific) following the manufacturer’s recommendations. A full list of primers is provided in <u>Table S8</u>.

Protoplast generation and transformation were carried out as previously described (Földi et al., 2024)^31^. Briefly, wild-type *C. cinerea* was grown on YMG medium under light at 37 °C for 6 days to induce oidium formation. Oidia were collected by adding 5 mL distilled water and gently brushing the colony surface, then transferred to 50 mL YMG and incubated overnight at 37 °C with shaking (120 rpm). Germinated oidia were pelleted by centrifugation at 2,600 × g for 5 minutes, resuspended in 40 mL MM buffer containing 2% VinoTaste Pro (Novozymes), and digested at 37 °C for 3–4 hours with gentle shaking (100 rpm). Digestion was stopped by adding CaCl₂ to a final concentration of 25 mM. Protoplasts were filtered through a 40 µm cell strainer (VWR), pelleted at 640 × g for 10 minutes at 4 °C, and washed once with 5 mL cold MMC buffer. For each transformation, protoplasts were resuspended in 100 µL MMC buffer and stored on ice until further use.

Protoplast transformation was carried out using a modified PEG/CaCl₂-based method adapted from Dörnte and Kües (2012)^90^. For each 100 µL protoplast suspension (∼10⁶–10⁷ cells), the transformation mix included ∼1 µg of repair template DNA, 15 µL of pre-assembled RNP complex, and 25 µL of PEG/CaCl₂ solution. To enhance membrane permeability and promote RNP uptake, Triton X-100 was added to the protoplast suspension at a final concentration of 0.006%^93^. The transformation mix was incubated on ice for 30 minutes, followed by a heat shock step in which 500 µL of PEG/CaCl₂ was added and the mixture was left at room temperature for 10 minutes. Transformed protoplasts were gently resuspended in 5 mL MM buffer and combined with 35 mL of molten top agar (<40 °C). Aliquots (10 mL) of this mixture were overlaid onto four regeneration plates (MM medium without PABA). Plates were incubated at 37 °C, and colonies began to appear within 3–4 days. Individual colonies were subsequently transferred to selective MM plates (lacking PABA) for downstream screening and analysis.

Gene disruption was confirmed using heat-assisted colony PCR, following the protocol of Dörnte and Kües (2013)^94^. Briefly, a small amount of mycelium was suspended in 100 µL sterile distilled water, boiled for 1 minute, vortexed, and boiled again for 1 minute. Samples were stored at −20 °C for at least 10 minutes before use. After thawing, they were centrifuged and the supernatant served as the PCR template. Reactions were performed using Phire Green Master Mix (Thermo Scientific) with primers specific to each construct, following the manufacturer’s instructions. For PCR verification of homologous recombination, we used external checking primers located outside the homologous arms of the repair template (‘checking’) together with primers specific to the selection marker *pab1* (‘PABA’). All primers are listed in <u>Table S”Primers”</u>. To isolate genetically homogeneous strains, transformants were first cultured on YMG medium at 37 °C under continuous light for 5 days. Oidia were then harvested from the mycelial surface using 0.01% Tween-20, diluted, and plated on minimal medium to obtain individual colonies.

### Phenotyping

To obtain fruiting bodies at comparable developmental stages, strains were inoculated into 250-mL beaker glasses containing 100 mL YMG medium with half the standard glucose concentration and solidified with 0.6% agar. Fruiting was induced by a 10-mL block of 1% water agar supplemented with 0.2 mM CuSO₄ and 0.2 mM MgSO₄ on the top of YMG^95^. The mouth of the beaker was covered loosely with a sterile tinfoil cap. After the inoculation of the strain, the culture was incubated at 28°C in a 12h light / 12h dark incubator for about 4-5 days.

Spore quantification was performed using 100 μg of mature cap tissue from wild-type (n = 5) and Δ*psr2* (n = 4) strains. Samples were suspended in 1 mL of 0.01% Tween-20, vortexed for 5 minutes, and filtered through a 40 μm cell strainer. Spore counts were determined using a Bürker chamber, and spore size was measured with Leica light microscopy software.

Sampling from the gills of young fruiting bodies was performed at the 12^th^ hour of the light cycle. For nuclear staining, central gill fragments were excised using a scalpel and incubated in darkness for 15 minutes in 100 μL of Hoechst 33342 solution (10 μg/mL; Thermo Scientific). Following incubation, samples were washed twice with 200 μL phosphate-buffered saline (PBS). Fluorescence imaging was carried out using a Zeiss LSM 800 confocal microscope. As the excitation maximum of Hoechst 33342 is at 361 nm and the emission maximum at 497 nm, samples were excited and detected in the ultraviolet range. The obtained fluorescence and brightfield images were overlaid using the ImageJ/Fiji software (version 1.54)^96^. Fluorescent images acquired at different focal planes were color-coded according to optical sections, and brightfield images from multiple planes were merged while retaining maximum intensity.

## Acknowledgements

We appreciate the support of the Doctoral Student Scholarship Program of the Co-operative Doctoral Program (KDP-17-4/PALY-2021) of the Ministry of Innovation and Technology financed from the National Research, Development and Innovation Fund. The Authors also acknowledge support by the National Research Development and Innovation Office (Grant No. OTKA 142188), the European Research Council (grant no. 101086900 to L.G.N.). DASz appreciates support of the National Academy of Scientist Education Program of the National Biomedical Foundation under the sponsorship of the Hungarian Ministry of Culture and Innovation. Z.M. was supported by the Janos Bolyai Research Scholarship of the Hungarian Academy of Sciences (BO/00269/24/8).

## Author Contributions

C.F., L. G. and L.G.N. conceived the study. C.F. generated mutants, performed the experiments, analyzed the data, and prepared figures. E.A. and Z.L. helped with fluorescent microscopy. Z.H., Z.M., X.L., A.Cs., D.A.Sz. and C.F. did the time-lapse videography and photography of fruiting. Z.M., C.F., B.B., and B.H. analyzed the bioinformatic data, C.F., L.G.N., and Z.M. wrote the paper. All authors have read and commented on the manuscript.

